# Deciphering spatial domains from spatially resolved transcriptomics with adaptive graph attention auto-encoder

**DOI:** 10.1101/2021.08.21.457240

**Authors:** Kangning Dong, Shihua Zhang

**Affiliations:** NCMIS, CEMS, RCSDS, Academy of Mathematics and Systems Science, Chinese Academy of Sciences, Beijing 100190, China; School of Mathematical Sciences, University of Chinese Academy of Sciences, Beijing 100049, China; Center for Excellence in Animal Evolution and Genetics, Chinese Academy of Sciences, Kunming 650223, China; Key Laboratory of Systems Biology, Hangzhou Institute for Advanced Study, University of Chinese Academy of Sciences, Chinese Academy of Sciences, Hangzhou 310024, China

## Abstract

Recent advances in spatially resolved transcriptomics have enabled comprehensive measurements of gene expression patterns while retaining the spatial context of the tissue microenvironment. Deciphering the spatial context of spots in a tissue needs to use their spatial information carefully. To this end, we developed a graph attention auto-encoder framework STAGATE to accurately identify spatial domains by learning low-dimensional latent embeddings via integrating spatial information and gene expression profiles. To better characterize the spatial similarity at the boundary of spatial domains, STAGATE adopts an attention mechanism to adaptively learn the similarity of neighboring spots, and an optional cell type-aware module through integrating the pre-clustering of gene expressions. We validated STAGATE on diverse spatial transcriptomics datasets generated by different platforms with different spatial resolutions. STAGATE could substantially improve the identification accuracy of spatial domains, and denoise the data while preserving spatial expression patterns. Importantly, STAGATE could be extended to multiple consecutive sections to reduce batch effects between sections and extracting three-dimensional (3D) expression domains from the reconstructed 3D tissue effectively.

## Introduction

The functions of complex tissues are fundamentally related to the spatial context of different cell types^1^. The relative locations of transcriptional expressions in tissue are critical for understanding its biological functions and describing interactive biological networks^2^. Breakthrough technologies for spatially resolved transcriptomics (STs), such as 10x Visium^3^, Slide-seq^4, 5^, and Stereo-seq^6^, have enabled genome-wide profiling of gene expressions in captured locations (referred to as spots) at a resolution of several cells or even subcellular levels.

Deciphering spatial domains (i.e., regions with similar spatial expression patterns) is one of the great challenges from STs. Most existing clustering methods do not efficiently use the available spatial information. These non-spatial methods can be roughly divided into two categories. The first category uses traditional clustering methods such as k-means and Louvain algorithm^7^. These methods are limited to the small number of spots or the sparsity according to the different resolutions of ST technologies, and clustering results may be discontinuous in the tissue section. The second category utilizes the cell type signatures defined by single-cell RNA-seq to deconvolute the spots^8, 9^. While these integration methods are appealing, as the spatial resolution improving, they are not applicable to ST data at a resolution of cellular or subcellular levels.

Some recent algorithms adapt the clustering methods by considering the similarity between adjacent spots to better account for the spatial dependency of gene expressions^10-12^. These methods show significant improvements in identifying spatial domains of sections from brain and cancer tissues. For example, BayesSpace is a Bayesian statistical method that encourages neighboring spots to belong to the same cluster by introducing spatial neighbor structure into the prior^10^. stLearn defines the morphological distance based on features extracted from a histology image and utilizes such distances as well as spatial neighbor structure to smooth gene expressions^11^. SEDR employs a deep auto-encoder network for learning gene representations and uses a variational graph auto-encoder to simultaneously embed spatial information^12^. Although these methods consider the spatial structure of STs, the similarity of neighboring spots defined by them is pre-defined before training and cannot be learned adaptively. Moreover, these methods do not consider the spatial similarity of spots at the boundary of spatial domains in more detail and do not well integrate spatial information to impute and denoise gene expressions. More importantly, these approaches cannot be applied to multiple consecutive sections to reconstruct a three-dimensional (3D) ST model and extract 3D expression domains **(Supplementary Table S1)**.

To this end, we developed a graph attention auto-encoder framework STAGATE to accurately identify spatial domains by learning low-dimensional latent embeddings via integrating spatial information and gene expression profiles. Extensive tests and comparison with existing methods on ST data generated by different platforms (e.g., 10x Visium, Slide-seq, and Stereo-seq) as benchmarks demonstrated its superiorities for downstream analysis tasks such as spatial domain identification, visualization, spatial trajectory inference, data denoising, and 3D expression domain extraction.

## Results

### Overview of STAGATE

STAGATE first constructs the spatial neighbor network (SNN) based on the spatial locations, and optionally introduces the cell type-aware SNN by pruning the SNN based on the pre-clustering of gene expressions **(Fig. 1)**. The gene expression pre-clustering can effectively identify regions containing distinct cell types, thus this cell type-aware SNN can help to better characterize the spatial similarity at the boundary of these distinct spatial domains for ST data with low spatial resolutions, such as 10x Visium **(Materials and Methods)**.

**Figure 1.**
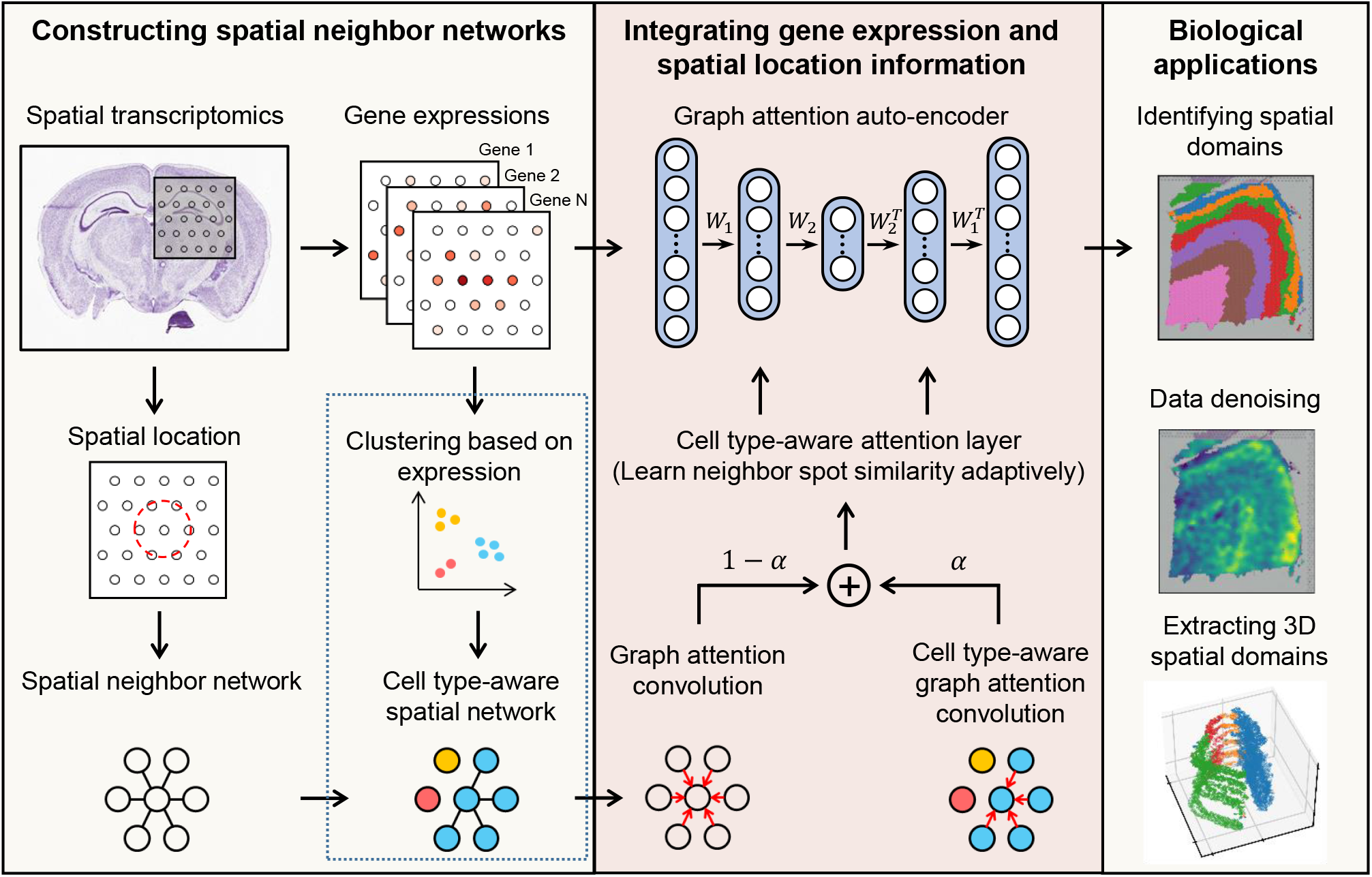
Overview of STAGATE. STAGATE first constructs a spatial neighbor network (SNN) based on a pre-defined radius, and another optional one in the dashed box for 10x Visium data by pruning it according to the pre-clustering of gene expressions to better characterize the spatial similarity at the boundary of spatial domains. STAGATE further learns low-dimensional latent representations with both spatial information and gene expressions via a graph attention auto-encoder. The input of the auto-encoder is the normalized expression matrix, and the graph attention layer is adopted in the middle of the encoder and decoder. The output of STAGATE can be applied for identifying spatial domains, data denoising, and extracting 3D spatial domains.

Then STAGATE learns low-dimensional latent embeddings with both spatial information and gene expressions via a graph attention auto-encoder^13^ **(Fig. 1)**. The normalized expression of each spot is first transformed into a *d*-dimensional latent embedding by an encoder and then reversed back into a reconstructed expression profile via a decoder. Unlike the classic auto-encoder, STAGATE adopts an attention mechanism in the middle layer of the encoder and decoder. It adaptively learns the edge weights of SNNs (i.e., the similarity between neighboring spots) and further uses them to update the spot representation by collectively aggregating information from its neighbors. Finally, the latent embeddings are used to visualize the data with UMAP^14^ and identify spatial domains with various clustering algorithms, such as mclust^15^ and Louvain^7^ **(Fig. 1)**.

### STAGATE improves the identification of known layers on the human dorsolateral prefrontal cortex dataset

To quantitatively evaluate the spatial clustering performance of STAGATE, we first applied it onto a 10x Visium dataset containing spatial expressions of 12 human dorsolateral prefrontal cortex (DLPFC) sections^16^. Maynard et al.^16^ has manually annotated DLPFC layers and white matter (WM) based on the morphological features and gene markers **(Fig. 2a)**. Considering it as the ground truth, we compared the clustering accuracy of STAGATE with the non-spatial clustering method implemented by SCANPY^17^ and three recently developed spatial clustering approaches (BayesSpace^10^, stLearn^11^, and SEDR^12^) in terms of adjusted rand index (ARI) **(Supplementary Notes)**.

**Figure 2.**
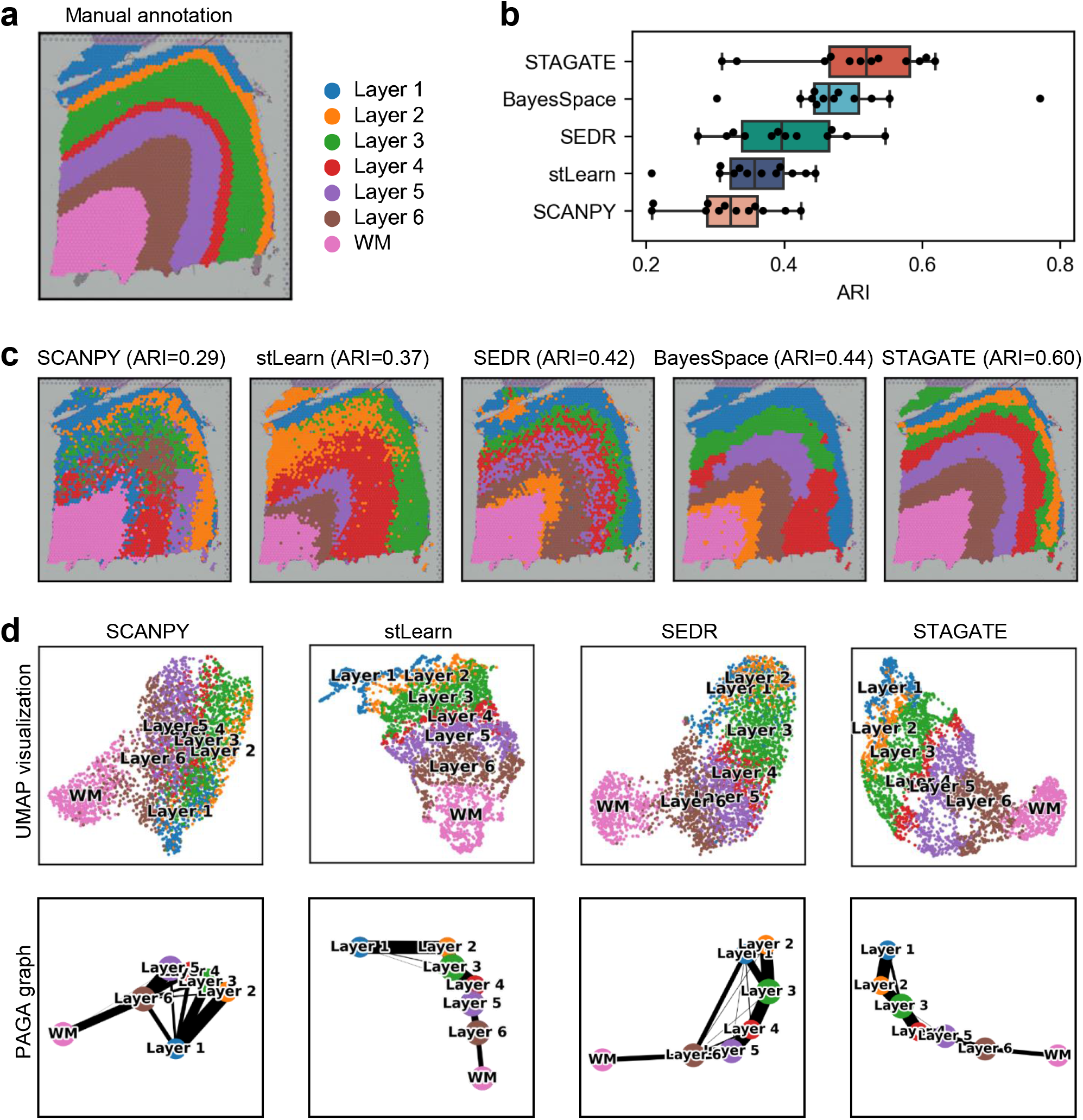
STAGATE improves the identification of layer structures in the human dorsolateral prefrontal cortex (DLPFC) tissue. **a**, Ground-truth segmentation of cortical layers and white matter (WM) in the DLPFC section 151676. **b**, Boxplot of clustering accuracy in all 12 sections of the DLPFC dataset in terms of adjusted rand index (ARI) scores for five methods. **c**, Cluster assignments generated by SCANPY, stLearn, SEDR, BayesSpace, and STAGATE in the DLPFC section 151676. **d**, UMAP visualizations and PAGA graphs generated by SCANPY, stLearn, SEDR, and STAGATE embeddings respectively in the DLPFC section 151676. As an end-to-end clustering approach, BayesSpace cannot be visualized using UMAP and PAGA.

STAGATE could effectively identify the expected cortical layer structures and achieve significant improvement compared to other methods **(Fig. 2b and Supplementary Fig. S1)**. For example, in the DLPFC section 151676, STAGATE delineated the layer borders clearly and achieved the best clustering accuracy (ARI=0.60) **(Fig. 2c)**. For comparison, the clustering assignment of the non-spatial approach SCANPY could roughly follow the expected layer pattern in this section, but the boundary of its clusters was discontinuous with many outliers, which impaired its clustering accuracy. Interestingly, the performance of algorithms leveraging the spatial information (STAGATE, BayesSpace, SEDR, and stLearn) are significantly better than the non-spatial clustering method SCANPY. These results demonstrated the superiority of STAGATE at spatial domain identification and the necessity of its usage of spatial information.

The integration of spatial information enables STAGATE to reveal the distance between spatial domains and depict the spatial trajectory in a UMAP plot^14^. For example, in the DLPFC section 151676, cortical layers were well-organized and showed consistent spatial trajectories (from layer 1 to layer 6 and white matter) in the UMAP plots generated by the STAGATE embeddings **(Fig. 2d and Supplementary Fig. S2)**. This result is consistent with the functional similarity between adjacent cortical layers as well as the chronological order^18^. By contrast, in the UMAP plots of SCANPY embeddings, spots belonging to distinct layers were not separated clearly. As for the other two spatial clustering methods, stLearn did not distinguish WM and cortex layers clearly, and SEDR mixed the spots of layer 1 and layer 6. We further confirmed the inferred trajectory using a trajectory inference algorithm named PAGA^19^ **(Fig. 2d)**. The PAGA graphs of both STAGATE and stLearn embeddings showed a nearly linear development trajectory from layer 1 to layer 6 as well as the similarity between adjacent layers, while the PAGA results of both SCANPY and SEDR embeddings were mixed.

### STAGATE enables the identification of tissue structures from ST data of different spatial resolutions

We further tested whether STAGATE can be applied to ST data of different spatial resolutions. We first applied STAGATE onto a Slide-seqV2 dataset with 10μm spatial resolution from mouse hippocampus^5^. Compared to the 10x Visium platform with a resolution of 55μm, Slide-seqV2 can profile spatial expressions at a resolution of cellular levels with more spots (>10,000 per section) but less sequence depth per spot **(Supplementary Table S2)**. As expected, using the Louvain clustering algorithm with the same parameter, STAGATE can well characterize the tissue structures and uncover the spatial domains, while the clusters identified by SCANPY and SEDR lack clear spatial separation **(Fig. 3a and Supplementary Fig. S3)**. For example, STAGATE depicted a clear “cord-like” structure as well as an “arrow-like” structure in the hippocampal region and identified four spatial domains of it. This result is consistent with the annotation of hippocampus structures from the Allen Reference Atlas^20^ **(Fig. 3b)**. Specifically, the “cord-like” structure corresponds to the pyramidal layer of Ammon’s horn, which can be further separated into fields CA1, CA2, and CA3 (i.e., CA1sp, CA2sp, and CA3sp), and the “arrow-like” structure corresponds to the granule cell layer of the dentate gyrus (i.e., DG-sg). Although the CA2sp domain was not clustered separately due to the small spot number, it was separated in the UMAP plot of STAGATE embeddings **(Supplementary Fig. S4)**. Furthermore, the expressions of many known gene markers also verified the cluster partition of STAGATE **(Fig. 3c and Supplementary Fig. S5)**. For example, ITPKA and BCL11B showed differential expressions between domains of Ammon’s horn and are highly expressed at CA1sp as expected^21, 22^. The known molecular markers of hippocampal CA2 such as AMIGO2 and PCP4 were specifically expressed in the identified CA2sp domain^23^. In addition, LRRTM4 that has been found to mediate excitatory synapse development on dentate gyrus granule cells was specifically expressed at the identified DG-sg region^24^. Besides these known tissue structures, STAGATE also identified many well-separated spatial domains and revealed their spatial gene expression patterns via differential expression analysis **(Supplementary Fig. S5)**. For example, the domain within the hippocampus except for the “cord-like” and “arrow-like” structures (domain 2) exhibited strong expression of astrocytes gene markers DDN and CAMK2A^25^. The domain surrounding the hippocampal region (domain 7) expressed many oligodendrocytes-related gene markers such as TRF and MOBP^26^. Moreover, we also observed significant spatial expression patterns in the spatial domains 3 and 4 with the dominant expression of ENPP2 and NWD2 respectively. These results demonstrated that STAGATE can dissect spatial heterogeneity and further uncover spatial expression patterns. We also tested STAGATE on the mouse hippocampus section profiled by Slide-seq and 10x Visium technologies. As the initial version of Slide-seqV2, the transcript detection sensitivity of Slide-seq is relatively lower^4^ **(Fig. 3d)**. STAGATE depicted the known tissue structures well except CA2sp on the Slide-seq data **(Fig. 3e)** and 10x Visium data **(Fig. 3f)** respectively.

**Figure 3.**
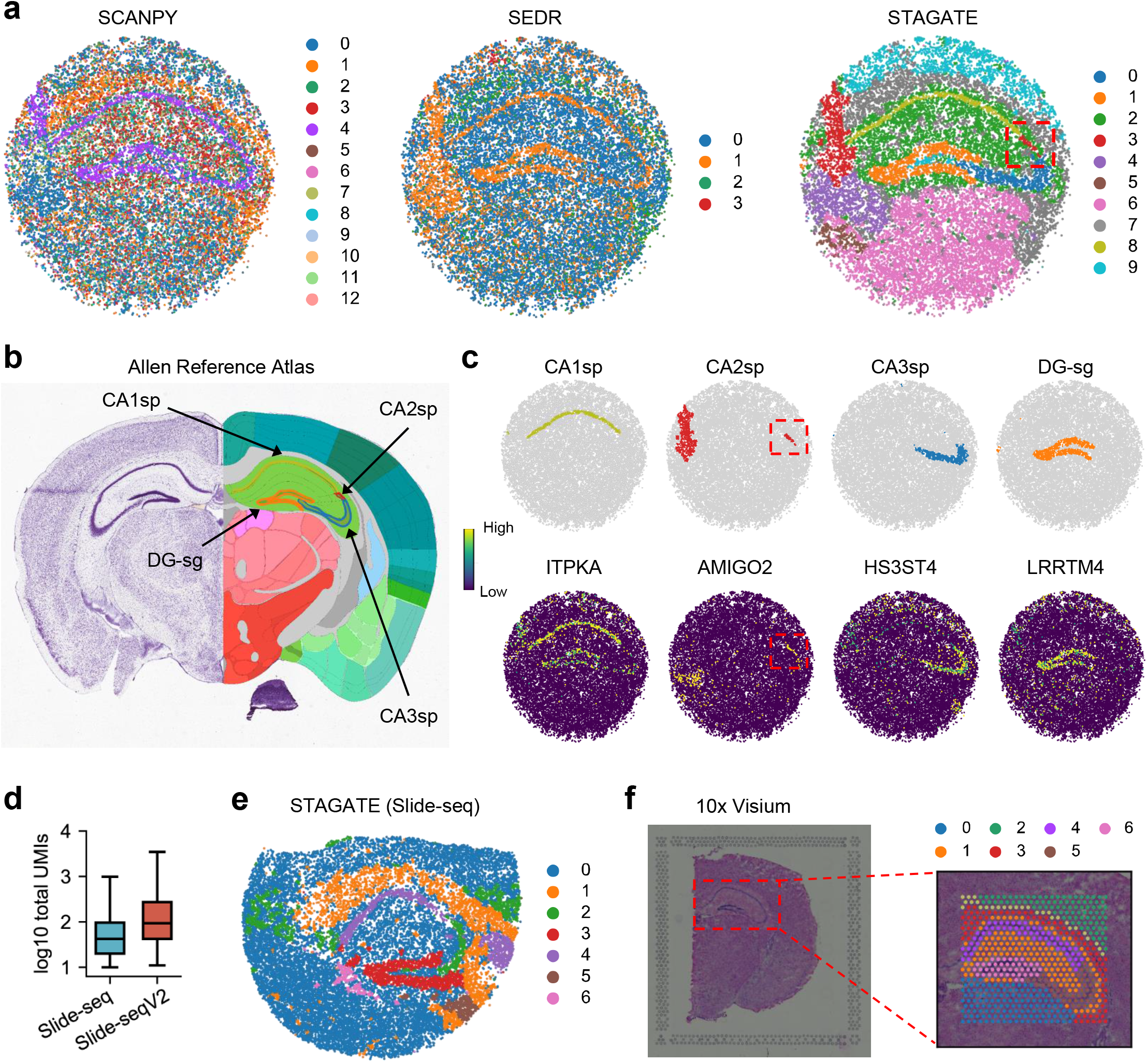
STAGATE improves the identification of known tissue structures in the mouse hippocampus tissue. **a**, Spatial domains generated by Louvain clustering with resolution=0.3 on the low-dimensional SCANPY, SEDR, and STAGATE embeddings in the Slide-seqV2 hippocampus section. **b**, The annotation of hippocampus structures from the Allen Reference Atlas of an adult mouse brain. **c**, Visualization of CA1sp, CA2sp, CA3sp, and DG-sg domains identified by STAGATE and the corresponding marker genes. Spatial domains were annotated by the structure annotation showed in the Allen Reference Atlas. **d**, Number of total UMIs per spot in the mouse hippocampus sections generated by Slide-seq and Slide-seqV2 respectively. **e** and **f**, Spatial domains generated by STAGATE on the hippocampus section profiled by Slide-seq (**e**) and 10x Visium (**f**) technologies respectively.

We also validated the performance of STAGATE for identifying tissue structures on the mouse olfactory bulb, a widely used model tissue with the laminar organization. We first tested STAGATE on a ST dataset generated by Stereo-seq from mouse olfactory bulb tissues^6^. Stereo-seq is a newly emerging spatial omics technology that can achieve the subcellular spatial resolution by DNA nanoball patterned array chips. The data used here were binned into a resolution of cellular levels (∼14μm)^6^. Fu et al.^12^ has annotated the laminar organization of coronal mouse olfactory bulb in the DAPI-stained image, containing the rostral migratory stream (RMS), granule cell layer (GCL), internal plexiform layer (IPL), mitral cell layer (MCL), external plexiform layer (EPL) and olfactory nerve layer (ONL) **(Fig. 4a)**. Compared to the clusters identified by SCANPY, those identified using both STAGATE and SEDR embeddings better reflected the laminar organization and well corresponded to the annotated layers **(Fig. 4b and Supplementary Fig. S6)**. Importantly, STAGATE recognized the narrow tissue structure MCL clearly, which was validated by the expression of mitral cell marker GABRA1^27^ **(Supplementary Fig. S7)**.

**Figure 4.**
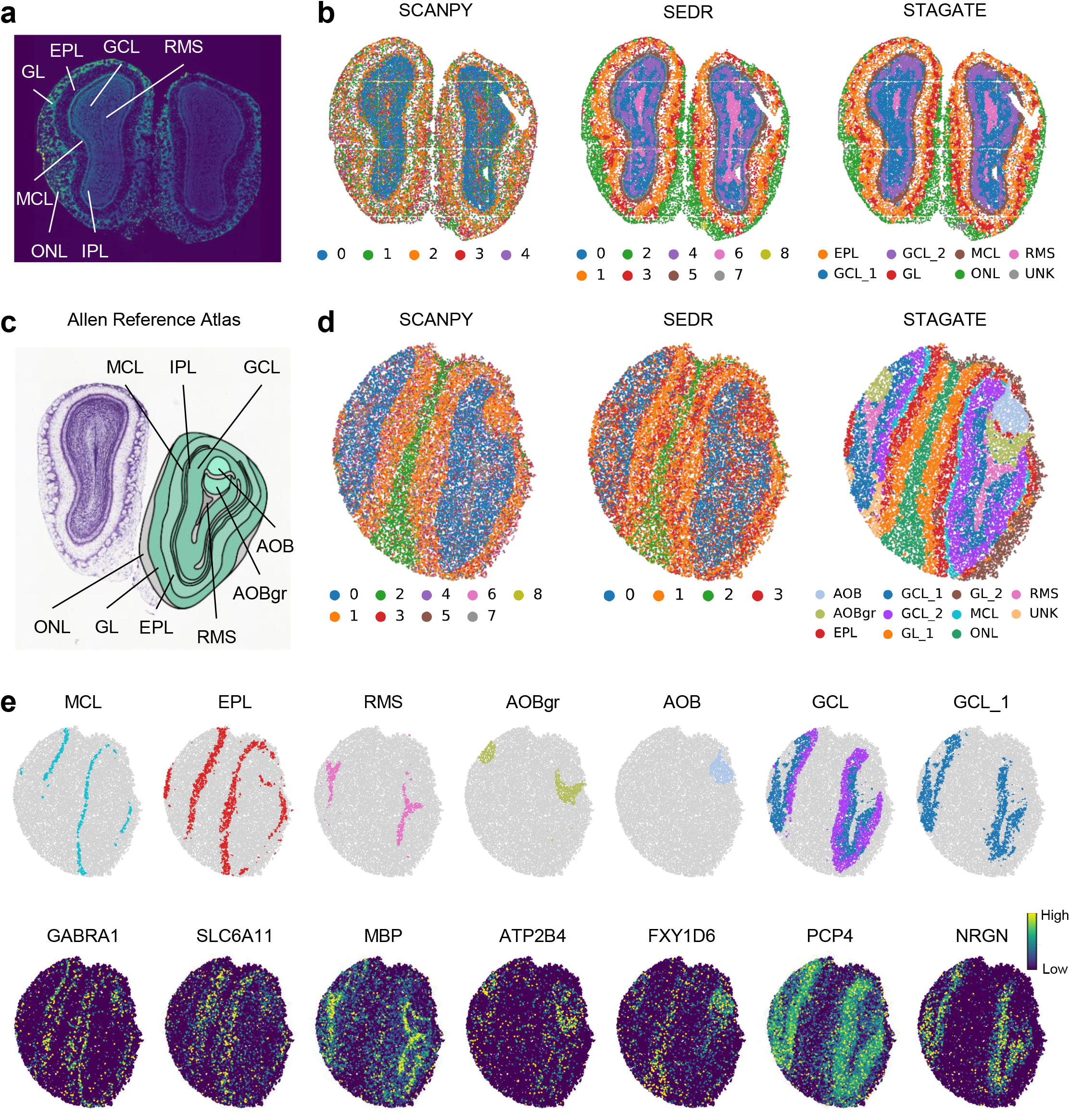
STAGATE identifies the laminar organization in the mouse olfactory bulb tissue sections profiled by Stereo-seq and Slide-seqV2 respectively. **a**, Laminar organization of mouse olfactory bulb annotated in the DAPI-stained image generated by Stereo-seq. **b**, Spatial domains generated by Louvain clustering with resolution=0.8 on the low-dimensional SCANPY, SEDR, and STAGATE embeddings in the Stereo-seq mouse olfactory bulb tissue section. **c**, Laminar organization of mouse olfactory bulb annotated by the Allen Reference Atlas. **d**, Spatial domains generated by Louvain clustering with resolution=0.5 on the low-dimensional SCANPY, SEDR, and STAGATE embeddings in the Slide-seqV2 mouse olfactory bulb tissue section. **e**, Visualization of spatial domains identified by STAGATE and the corresponding marker genes. Spatial domains were annotated by the laminar organization showed in the Allen Reference Atlas.

We also applied STAGATE onto a mouse olfactory bulb section profiled by Slide-seqV2^5^ and found that the spatial domains identified by STAGATE were well consistent with the annotation of coronal mouse olfactory bulb from Allen Reference Atlas^20^ **(Fig. 4c)**. Specifically, compared to clusters produced by SCANPY and SEDR, STAGATE identified two spatial domains corresponding to the accessory olfactory bulb (AOB) and the granular layer of the accessory olfactory bulb (AOBgr) respectively **(Fig. 4d and Supplementary Fig. S8)**. These spatial domains uncovered by STAGATE were clearly supported by known gene markers **(Fig. 4e)**. For example, FXYD6 showed strong expressions on the identified AOB domain, which is consistent with its immunohistochemistry experiment^28^. The granular cell marker ATP2B4^29^ showed strong expressions on the identified AOBgr domain. The narrow MCL structure with the dominant expression of mitral cell marker GABRA1^27^ was also identified by STAGATE. In addition, STAGATE identified a spatial subpopulation of GCL named GCL_1 with the dominant expression of NRGN. NRGN is a well-documented schizophrenia risk gene^30^, implying that this domain is related to cognition function. Moreover, we found that STAGATE delineated the spatial trajectory among the mouse olfactory bulb (from AOBgr to RMS to ONL) in the UMAP plots as well as the PAGA graphs **(Supplementary Fig. S9)**. Collectively, these results illustrated the ability of STAGATE to identify tissue structures and reveal their organization from ST data of different spatial resolutions.

### Attention mechanism and cell type-aware module help to better characterize the similarity between neighboring spots

Next, we tested whether STAGATE could provide insights into sections including more biologically complex tissues, such as the whole brain. We applied STAGATE onto a 10x Visium dataset, which profiled the spatial expressions of a coronal mouse brain section **(Fig. 5a)**. We found that the clustering results identified by SCANPY roughly divided the tissue structures containing different cell types while lacking the identification of small spatial domains **(Fig. 5b and Supplementary Fig. S10)**. For example, the clustering assignment of SCANPY failed to identify the “cord-like” structure -- Ammon’s horn and the “arrow-like” structure -- dentate gyrus within the hippocampus. Moreover, SEDR only smoothed the domain border, but cannot depict the small spatial domains either **(Fig. 5b)**. The direct application of STAGATE brought some improvements in spatial domain identification **(Fig. 5b)**. Specifically, in the hippocampal region, STAGATE without cell type-aware module identified the field CA1 (domain 17) and CA3 (domain 19) of Ammon’s horn, but did not depict the dentate gyrus structure.

**Figure 5.**
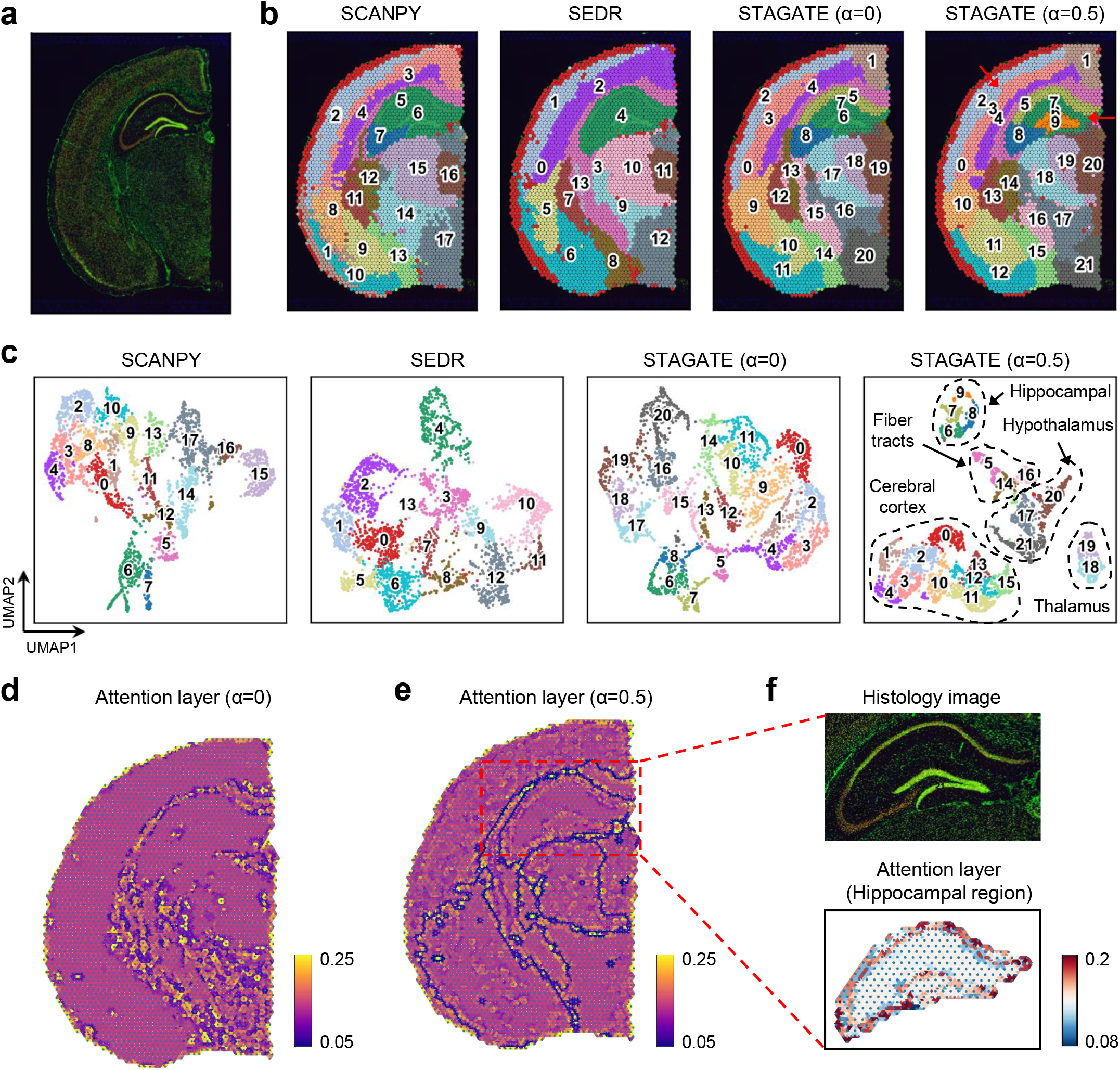
STAGATE reveals spatial domains in adult mouse brain section profiled by 10x Visium. **a**, Immunofluorescent imaging of the tissue section stained with DAPI and Anti-NeuN. **b**, Spatial domains generated by Louvain clustering with resolution=1 on the low-dimensional embeddings of SCANPY, SEDR, STAGATE, and STAGATE with cell type-aware module. **c**, UMAP visualizations of the low-dimensional embeddings of SCANPY, SEDR, STAGATE, and STAGATE with cell type-aware module respectively. **d** and **e**, Visualizations of the attention layer of STAGATE without (**d**) or with (**e**) cell type-aware module. The nodes of the attention layer are arranged according to the spatial position of spots. The edges of the attention layer are colored by corresponding weights. **f**, Zoomed-in views of immunofluorescent imaging of the hippocampus region and the visualization of attention layer in (**e**).

For ST data containing heterogeneous cell types with low spatial resolution, STAGATE with the cell type-aware module could better learn the spatial similarity **(Fig. 1; Materials and Methods)**. Specifically, the pre-clustering process is based on the Louvain algorithm with a small resolution parameter (set as 0.2 by default) **(Supplementary Fig. S10b)**. As expected, the usage of the cell type-aware module aided in the identification of spatial domains **(Fig. 5b)**. STAGATE identified the Ammon’s horn as well as the dentate gyrus structure in the hippocampus, and further depicted the spatial domains CA1 (domain 17) and CA3 (domain 20) of the Ammon’s horn. In addition, STAGATE better depicted the layer structures of the cortex region (domain 0, 4, and 12). Notably, we found that the cell type-aware module also significantly improved the separation of tissue structures in the UMAP plot, while those of SEDR and STAGATE without cell type-aware module were more like a smooth version of the non-spatial method SCANPY **(Fig. 5c)**.

We further evaluated whether the usage of attention mechanism indeed contributed to better characterizing the heterogeneous similarity between neighboring spots. We visualized the attention layer by arranging the nodes according to their spatial locations and coloring the edges by their weights, and found that using the attention mechanism alone could delineate the boundaries of main tissue structures such as the cortex, hippocampus, and midbrain **(Fig. 5d)**. Combining the attention mechanism and the cell type-aware module enhanced the delineation of structure boundaries, and further revealed the spatial similarity within small spatial domains **(Fig. 5e)**. For example, in the hippocampal region, STAGATE adaptively learned the spatial similarity within the Ammon’s horn as well as the dentate gyrus structure **(Fig. 5f)**. Collectively, these results indicated the importance of the attention mechanism and cell type-aware module for depicting the similarity between neighboring spots.

### STAGATE denoises gene expressions for better characterizing spatial expression patterns

STAGATE could denoise and impute gene expressions. We adopted STAGATE to reduce noises in the DLPFC dataset to better show the spatial pattern of genes. We compared the expressions of six layer-marker genes of the raw data to those denoised ones by STAGATE in the DLPFC section 151676 **(Fig. 6a)**. As expected, the denoised ones by STAGATE exhibited the laminar enrichment of these layer-marker genes clearly. For example, after denoising, the ATP2B4 gene showed differential expressions in layers 2 and 6, which is consistent with previously reported results^16, 31^, while its raw spatial expression is completely messy. We validated the laminar enrichment showed by STAGATE against publicly available in situ hybridization (ISH) data from the Allen Human Brain Atlas^20^ **(Fig. 6b)**. Moreover, a comparison of the raw expressions and the denoised ones by STAGATE using violin plots demonstrated that STAGATE enhanced the spatial patterns of layer-marker genes **(Fig. 6c, d)**. Notably, STAGATE obtained similar performance on the DLPFC section 151507 **(Supplementary Fig. S11)**. Collectively, these results demonstrated the ability of STAGATE to reduce noises and enhance spatial expression patterns. In addition, we also compared the imputation performance of STAGATE with four widely used single-cell RNA-seq imputation algorithms in terms of downsampling experiments, and showed its superior in both imputation efficiency and preservation of spatial expression patterns **(Supplementary Notes; Supplementary Fig. S12)**.

**Figure 6.**
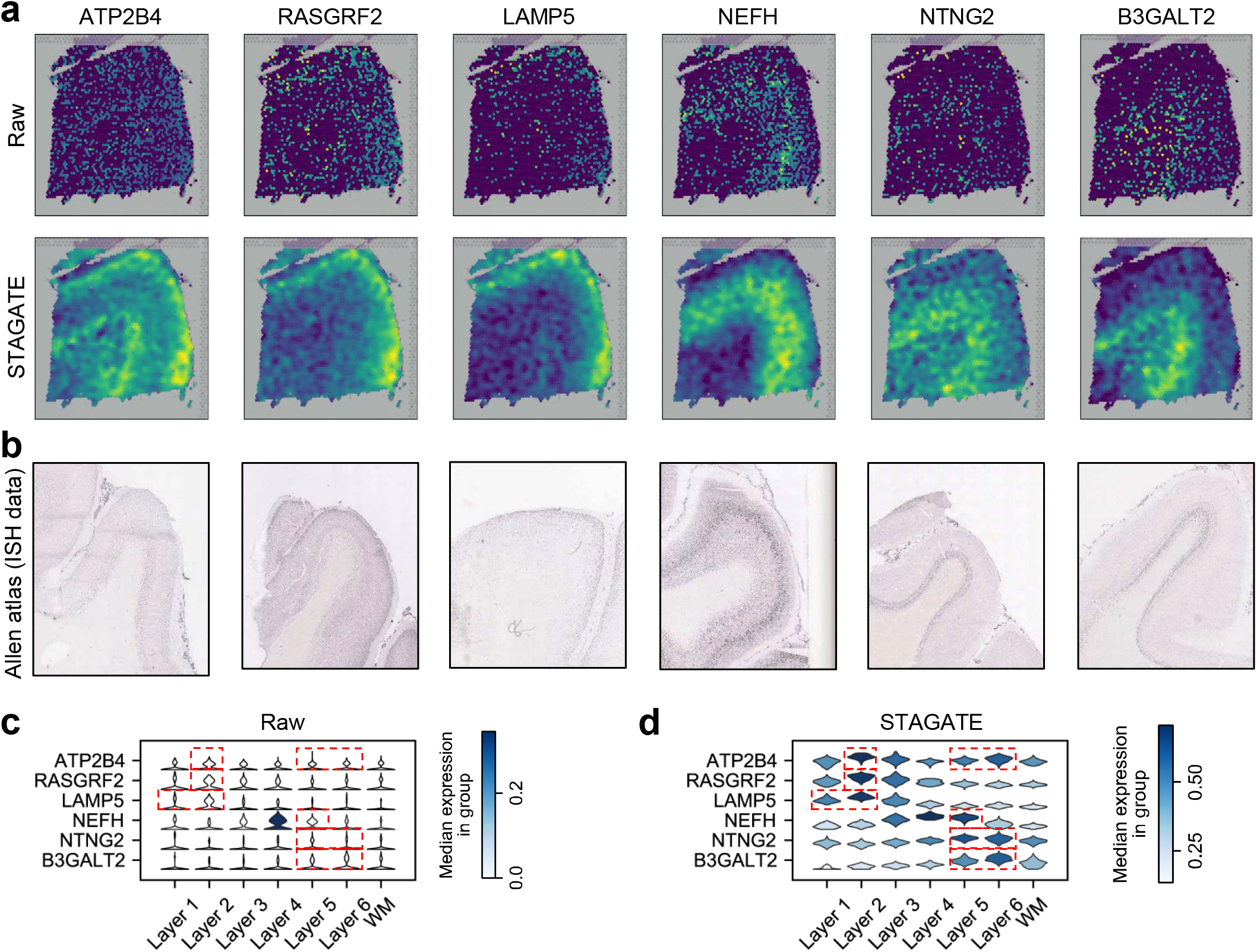
STAGATE enhances the spatial patterns of layer-marker genes in the DLPFC dataset. **a**, Visualizations of the raw spatial expressions and STAGATE denoised ones of six layer-marker genes in the DLPFC section 151676. **b**, ISH images from visual cortex (ATP2B4, RASGRF2, NEFH, NTNG2, and B3GALT2) or temporal cortex (LAMP5) of the adult human brain from the Allen Human Brain Atlas. **c**, Violin plots of the raw expression of layer-marker genes. **d**, Violin plots of the STAGATE denoised expressions of layer-marker genes. The cortical layer corresponding to the layer-marker gene is marked with a red box.

### The usage of 3D SNN leads to better extraction of 3D spatial patterns

We applied STAGATE onto a pseudo-3D ST data constructed by aligning the spots of the “cord-like” structure in seven hippocampus sections profiled by Slide-seq **(Fig. 7a; Supplementary Table S3)**. We extended STAGATE for 3D spatial domain identification by simultaneously considering the 2D SNN within each section and neighboring spots between adjacent sections **(Fig. 7b; Materials and Methods)**. Due to the data sparsity, the clustering results generated using SCANPY were mixed **(Supplementary Fig. S13)**. When only adopting the 2D SNN, STAGATE failed to identify the CA2sp domain due to the batch effects between sections **(Fig. 7c, d)**. After adding neighboring edges between adjacent sections, STAGATE depicted the known tissue structures clearly, and spots tend to cluster by their spatial structures rather than by section IDs in the UMAP plot **(Fig. 7e, f)**. We verified the tissue structures identified based on STAGATE by the known marker genes, including ITPKA^21^, BCL11B^22^, AMIGO2^23^, and LRRTM4^24^ **(Supplementary Fig. S13d)**. These results illustrated that STAGATE could help to reconstruct 3D tissue models and accurately extract 3D expression patterns by incorporating 3D spatial information.

**Figure 7.**
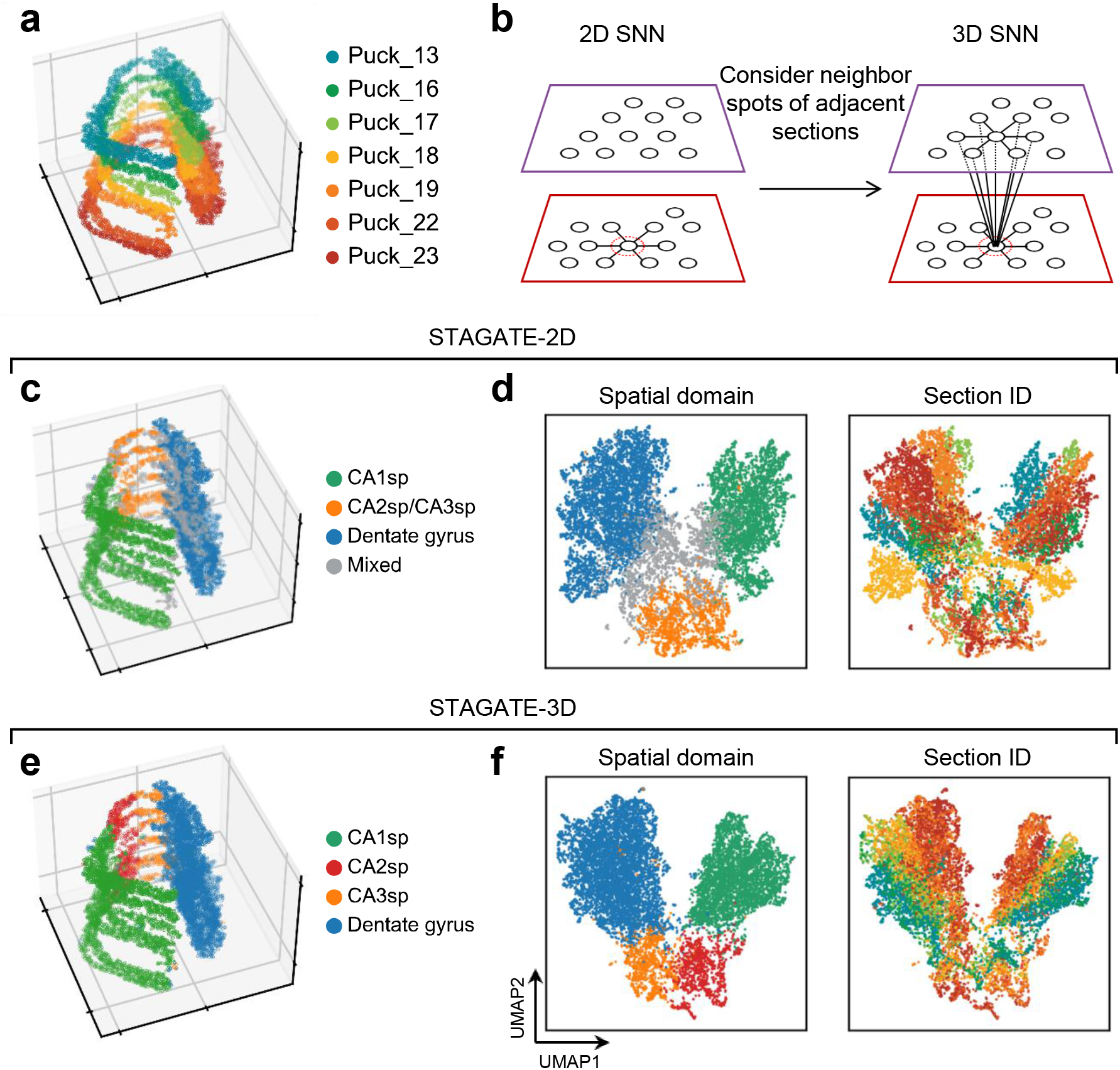
STAGATE can alleviate the batch effect between consecutive sections by incorporating a 3D spatial network. **a**, Visualization of the 3D hippocampal volume stacked by seven aligned consecutive sections profiled by Slide-seq. **b**, The 3D SNN is a combination of the 2D SNN within each section and the spatial network between consecutive sections. **c**, Cluster assignments generated by STAGATE-2D with the 2D SNN. **d**, The UMAP plots generated by STAGATE-2D embeddings. The spots are colored by the identified spatial domains (left) and the section IDs (right) respectively. **e**, Cluster assignments generated by STAGATE-3D with the 3D SNN. **f**, The UMAP plots generated by STAGATE-3D embeddings. The spots are colored by the identified spatial domains (left) and the section IDs (right) respectively.

## Discussion

Accurate identification of spatial domains and further extraction of spatially expressed genes are essential for understanding tissue organization and biological functions. Here, we developed a fast and user-friendly spatial domain identification method STAGATE, which can be seamlessly integrated into the standard analysis workflow by taking the “anndata” object of SCANPY package^17^ as inputs. STAGATE transforms spatial location information into SNNs and further adopts a graph attention auto-encoder to integrate SNNs and expression profiles. We tested the performance of STAGATE on diverse ST data generated by different platforms with different spatial resolutions. We found that STAGATE accurately revealed the laminar organization of DLPFC and mouse olfactory bulb. Moreover, STAGATE identified the known tissue structures of the hippocampus clearly and uncovered spatial domains of it. We additionally demonstrated the ability of STAGATE in expression denoising by comparing it with the ISH images. Lastly, we illustrated the ability of STAGATE to alleviate batch effects between consecutive sections and extract 3D expression domains in a pseudo-3D ST model.

The success of STAGATE is mainly attributed to the usage of the graph attention mechanism for considering spatial neighbor information. However, the current STAGATE focuses on the integration of expression profiles and spatial information and does not leverage the histological images. Existing methods taking histological images as inputs, such as stLearn, did not achieve good performance in our comparison. stLearn employs a pre-trained neural network to extract features from images and further calculates the morphological distance by cosine distance. We believe this pre-defined approach does not take advantage of the flexibility of deep learning, and the attention mechanism can be extended to adaptively integrate the histological image features conveniently.

STAGATE can handle ST data of diverse spatial resolutions. Generally, STAGATE performs better for ST data of cellular or subcellular resolutions due to the high similarity between neighboring spots. For technologies with relatively low spatial resolution, we introduced the cell type-aware module to depict the heterogeneous spatial similarity. However, a potential limitation of STAGATE is that it treats neighboring spots from one section the same as those belonging to different sections. Future work may employ heterogeneous networks to better depict 3D tissue models.

With the increase of spatial resolution and data scale, the computational approach should meet the basic requirement of efficiency and scalability. We recorded the running time spent on real datasets by STAGATE **(Supplementary Fig. S14a)**. When dealing with the largest real dataset with more than 50k spots, STAGATE only costs about 40 minutes. We also benchmarked the running time and memory usage of STAGATE on simulated datasets of different scales where spots were arranged according to the location of 10x Visium chips. Numerical experiments showed that STAGATE was fast and only took less than 40 minutes with about 4GB GPU memory usage for dealing with the dataset with 50k spots **(Supplementary Fig. S14b)**. However, the GPU memory usage is nearly linearly correlated to the number of spots and could be a bottleneck restricting the application of STAGATE to massive datasets **(Supplementary Fig. S14c)**. Future work is expected to improve the scalability of STAGATE by introducing the subgraph-based training strategy.

Moreover, STAGATE enables the detection of spatially variable genes within spatial domains. Existing spatially variable gene identification algorithms such as SPARK-X^32^ do not consider the spatial domain information, which makes it difficult to identify space-specifically expressed genes within small tissue structures. To illustrate it, we compared differential expressed genes of STAGATE spatial domains with those of SPARK-X on the Slide-seqV2 dataset from mouse olfactory bulb tissue. Specifically, STAGATE identified 959 domain-specific genes, and SPARK-X searched 2,479 spatially variable genes with FDR<0.01. We found that many genes identified by SPARK-X did not show significant differences between spatial domains **(Supplementary Fig. S15a)**. Furthermore, the spatial autocorrelations measured by Moran’s I statistic were similar between the gene set identified by STAGATE and the first 1,000 genes of SPARK-X **(Supplementary Fig. S15b)**. The gene sets identified by these two methods have a great overlap, but SPARK-X ignores some specific genes of small tissue structures **(Supplementary Fig. S15c)**. For example, the mitral cell marker GABRA1 show significant enrichment in the MCL domain **(Fig. 4e; FDR=1e-34)**, but SPARK-X did not identify its spatial pattern **(FDR =0.018)**. Moreover, the NEFH gene also showed a strong expression in the MCL domain **(Supplementary Fig. S15d; FDR=1e-12)**, while SPARK-X ignored it **(FDR=1)**. We expect that STAGATE can facilitate the identification of tissue organization and the discovery of corresponding gene markers.

## Materials and Methods

### Data description

We applied STAGATE to ST datasets generated by different platforms including 10x Visium, Slide-seq, Slide-seqV2, and Stereo-seq (see **Supplementary Table S2** for details). Specifically, the DLPFC dataset includes 12 human DLPFC sections sampled from three individuals experiments^16^. The number of spots ranges from 3,498 to 4,789 for each section, and the original authors have manually annotated the area of DLPFC layers and white matter (WM). The Stereo-seq mouse olfactory bulb data has been binned into a resolution of cellular levels (∼14μm) and contains 19,109 spots^12^. The Slide-seqV2 mouse hippocampus data and mouse olfactory bulb data were profiled at a spatial resolution of 10μm, and contain 19,285 and 20,139 spots respectively^5^.

Moreover, seven ST data profiled by Slide-seq were used to reconstruct the 3D hippocampus model^4^ (**Supplementary Table S3**). To generate the 3D hippocampus model, we first extracted the “cord-like” structure and “arrow-like” structure based on the STAGATE embeddings in the entire section. Then sections are aligned using the Iterative Closest Point algorithm^33^ and manual fine-tuning.

We also downloaded publicly available ISH images and the annotation atlas images from the Allen Brain Atlas website^20^ (**Supplementary Table S4**).

### Data preprocessing

In all datasets, we first removed spots outside the main tissue area. Then raw gene expressions were log-transformed and normalized according to library size using SCANPY package^17^. Finally, the top 3,000 highly variable genes were selected as the inputs of STAGATE.

### Construction of SNN

To incorporate the similarity of neighboring spots of a given spot, STAGATE converts the spatial information into an undirected neighbor network according to a pre-defined radius *r*. Let **A** be the adjacency matrix of the SNN, then A_*ij*_ = 1 if and only if the Euclidean distance between spot *i* and spot *j* is less than *r* **(Fig. 1)**. Specifically, for 10x Visium data, we set the network to contain the six nearest neighbors for each spot. For other data, we empirically choose *r* so that each spot contains 6-15 neighbors on average. The statistics of the number of neighbors for all the experiments can be found in **Supplementary Fig. S16**. Self-loops are added for each spot.

### Construction of cell type-aware SNN (optional)

For ST data with relatively low spatial resolution, STAGATE adopts a cell type-aware module by pruning the SNN according to pre-clustering of gene expressions. Specifically, the pre-clustering of gene expressions is conducted by the Louvain algorithm with a small resolution value (set as 0.2 by default) on the PCA embeddings, and STAGATE prunes the edge if the spots of it belong to different clusters **(Fig. 1)**.

### Graph attention auto-encoder

The graph attention auto-encoder consists of three parts: encoder, decoder and graph attention layer.

#### Encoder

The encoder in our architecture takes the normalized gene expressions as input and generates the spot embedding by collectively aggregating information from its neighbors. Let *x*_*i*_ be the normalized expressions of spot *i* and *L* be the number of layer of the encoder. By treating expression profiles as initial spot embeddings (i.e.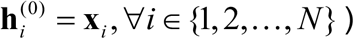), the *k*-th (*k ∈* {1, 2,.., *L −* 1}) encoder layer generates the embedding of spot *i* in layer *k* as follows:

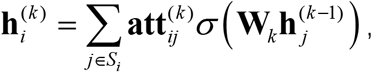

where **W**_*k*_ is the trainable weight matrix, σ is the nonlinear activation function, *𝒮*_*i*_ is the neighbor set of spot *i* in SNN (including spot *i* itself) and 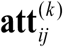 is the edge weight between spot *i* and spot *j* in the output of the *k*-th graph attention layer. The *L*-th encoder layer does not adopt the attention layer and is formulated as follows:

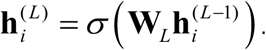

The output of encoder is considered as the final spot embedding.

#### Decoder

By contrast, the decoder reverses the latent embedding back into a reconstructed normalized expression profile. By treating the output of the encoder as the input of the decoder (i.e.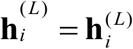), the *k*-th (*k ∈* {2,.., *L* −1, *L*}) decoder layer reconstructs the embedding of spot *i* in layer *k*-1 as follows:

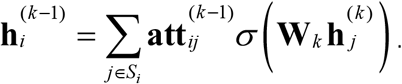

Similar to the encoder, the last layer of decoder is formulated as follows:

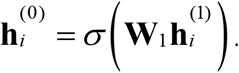

The output of decoder is considered as the reconstructed normalized expressions. To avoid overfitting, STAGATE sets 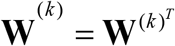 and **att**^(*k*)^ = **att**^(*k*)^ respectively.

#### Graph attention layer

To adaptively learn the similarity between neighboring spots, we employed a self-attention mechanism that has been widely used for graph neural networks^34^. We first described it in the context of using SNN alone. Briefly, the attention mechanism is a single-layer feedforward neural network with shared parameters among nodes, parametrized by a weight vector. In the *k*-th encoder layer, the edge weight from node *i* to its neighbor node *j* is computed as follows:

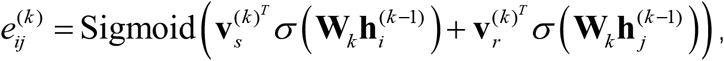

Where 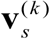 and 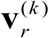 are the trainable weight vectors, and Sigmoid represents the sigmoid activation function.

To make the spatial similarity weights comparable, we normalized them by a softmax function as follows:

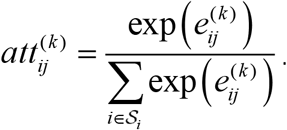

These learned weights are further used to update the latent embedding in the encoder and decoder.

In addition, when the cell type-aware module is used, STAGATE adopts self-attention mechanism for the two types of SNNs respectively. Let 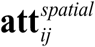 and 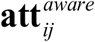 represent the learned spatial similarity based on SNN and the cell type-aware SNN respectively (the layer symbol is omitted here), and the spatial similarity finally adopted is the linear addition of them (α=0.5 by default):

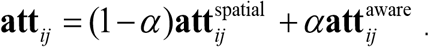

We discussed the selection of α on the 10x Visium coronal mouse brain data in **Supplementary Notes** and **Supplementary Fig. S17**.

#### Loss function

The objective of STAGATE is to minimize the reconstruction loss of normalized expressions as follows:

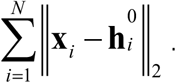

### The overall architecture of STAGATE

In all experiments, the encoder of STAGATE is set as a two-layer neural network (512-30) with the graph attention layer, and the decoder is set as the same number of layers as the encoder. Adam optimizer^35^ is used to minimize the reconstruction loss with an initial learning rate of 1e-4. The weight decay is set as 1e-4. The activation function is set as the exponential linear unit (ELU)^36^. The number of iterations is set as 500 by default, and 1,000 when using the cell type-aware module.

### Clustering

We used different strategies to identify spatial domains based on STAGATE embeddings. When the number of the label is known, we employ the mclust clustering algorithm^15^. For data without prior information, we use the Louvain algorithm implemented by SCANPY package^17^. The resolutions of the Louvain algorithm are manually selected. For fairness, we also displayed the results of the Louvain algorithm under different resolutions.

### Spatial trajectory inference

We employed the PAGA algorithm^19^ implemented in the SCANPY package^17^ to depict spatial trajectory. The PAGA graphs were visualized by the *scanpy*.*pl*.*paga_compare()* function.

### Identifying differentially expressed genes

We used the Wilcoxon test implemented in SCANPY package^17^ to identify differentially expressed genes for each spatial domain with a 1% FDR threshold (Benjamin-Hochberg adjustment).

### Identification of 3D spatial domains using STAGATE

All current ST technologies profile gene expression patterns in the context of 2D tissue sections, which limits the accurate depiction of 3D ST in the real world. A conventional solution is to reconstruct gene expressions in 3D space by stacking consecutive 2D sections^4, 37^. However, the batch effect between sections hinders the extraction of 3D spatial patterns. Here, we introduced a 3D SNN by incorporating the 2D SNN of each section and the SNN between adjacent sections to alleviate the batch effect between consecutive sections **(Fig. 7b)**. Specifically, the SNN between adjacent sections is constructed based on the aligned coordinates and a pre-defined radius. The key idea of the usage of 3D SNN is that the biological differences between consecutive sections should be continuous, so we can enhance the similarity between adjacent sections to eliminate the discontinuous independent technical noises.

## Data availability

All data analyzed in this paper are available in raw form from their original authors. Specifically, the DLPFC dataset is accessible within the *spatialLIBD* package (http://spatial.libd.org/spatialLIBD). The MouseBrain dataset is collected from the 10x Genomics website (https://support.10xgenomics.com/spatial-gene-expression/datasets). Slide-seqV2 datasets are available at https://singlecell.broadinstitute.org/single_cell/study/SCP815/highly-sensitive-spatial-transcriptomics-at-near-cellular-resolution-with-slide-seqv2#study-summary. Slide-seq datasets are available at https://portals.broadinstitute.org/single_cell/study/slide-seq-study. The processed Stereo-seq data from mouse olfactory bulb tissue is accessible on https://github.com/JinmiaoChenLab/SEDR_analyses.

## Acknowledgements

This work has been supported by the National Key Research and Development Program of China [2019YFA0709501]; the Strategic Priority Research Program of the Chinese Academy of Sciences (CAS) [XDPB17], the Key-Area Research and Development of Guangdong Province [2020B1111190001], the National Natural Science Foundation of China [61621003]; the National Ten Thousand Talent Program for Young Top-notch Talents, and the CAS Frontier Science Research Key Project for Top Young Scientist [QYZDB-SSW-SYS008].

## Author contributions

Shihua Zhang conceived and supervised the project. Kangning Dong designed, implemented, and validated STAGATE algorithm. Kangning Dong and Shihua Zhang wrote the manuscript. All authors read and approved the final manuscript.

